# Experimental evidence for delayed post-conflict management behaviour in wild dwarf mongooses

**DOI:** 10.1101/2021.05.02.442338

**Authors:** Amy Morris-Drake, Julie M. Kern, Andrew N. Radford

**Affiliations:** School of Biological Sciences, University of Bristol, 24 Tyndall Avenue, Bristol BS8 1TQ, UK; School of Environmental and Rural Science, University of New England, Armidale, New South Wales, 2351, Australia

## Abstract

In many species, within-group conflict leads to immediate avoidance of potential aggressors or increases in affiliation, but no studies have investigated delayed post-conflict management behaviour. Here, we experimentally test that possibility using wild dwarf mongooses (*Helogale parvula*). First, we used natural and playback-simulated foraging displacements to demonstrate that bystanders take notice of the vocalisations produced during such within-group conflict events. We then used another playback experiment to assess delayed effects of within-group conflict on grooming interactions. Overall, fewer individuals groomed on evenings following an afternoon of simulated conflict, but those that did groomed more than on control evenings. Subordinate bystanders groomed with the simulated aggressor significantly less, and groomed more with one another, on conflict compared to control evenings. Our study provides experimental evidence that dwarf mongooses acoustically obtain information about within-group contests (including protagonist identity), retain that information and use it to inform conflict-management decisions with a temporal delay.

## Introduction

Conflicts of interest are common in social species, with disagreements between group members arising over access to mates or food, synchronisation of group activities and the direction of travel (Aureli et al., 2002; Conradt and Roper, 2009; Hardy and Briffa, 2013). Within-group conflict, especially if it escalates to aggression, can be costly in terms of injury and mortality, time and energy expenditure, increased stress and disrupted social relationships (Aureli, 1997; Aureli et al., 2002; de Waal, 2000). Conflict-management strategies that minimise these costs, either by reducing the likelihood of aggressive escalation in the first place or by mitigating the consequences of such physical contests when they do arise, have therefore evolved in many species (Aureli et al., 2002; Aureli and de Waal, 2000). Much of the early work on post-conflict behaviour focussed on interactions between the protagonists (the aggressor and the victim): many studies have documented increases in affiliation between former opponents in the aftermath of a contest (reconciliation; Aureli et al., 2002; de Waal, 2000; de Waal and van Roosmalen, 1979), although there are also examples of victims avoiding aggressors (wariness; Benkada et al., 2020; Kutsukake and Clutton-Brock, 2008; Sommer et al., 2002). More recently, attention has shifted to the involvement of bystanders (contest nonparticipants) in post-conflict behaviour, particularly bystander-initiated affiliation with the victim as a means of avoiding redirected aggression (self-protection) or of providing substitute reconciliation or consolation (Fraser et al., 2009, 2008; Schino and Marini, 2012; Wittig and Boesch, 2010). There is also some evidence of bystander-initiated affiliation with the aggressor, which could function as appeasement to reduce the likelihood of redirected aggression (Cordoni and Palagi, 2015; Palagi et al., 2008; Pallante et al., 2018), and group-wide post-conflict affiliation among bystanders, perhaps to reduce conflict-induced anxiety (De Marco et al., 2010; Judge and Mullen, 2005). However, to the best of our knowledge, this research has focussed solely on interactions that occur in the immediate aftermath (usually within 10 minutes) of an aggressive contest; the possibility of delayed post-conflict management behaviour has not been explored.

There is increasing experimental evidence that nonhuman animals can remember past events and use information from them when making later social decisions (Carter and Wilkinson, 2013; Kern and Radford, 2018; Seyfarth and Cheney, 1984; Wittig et al., 2014). This includes conflict-management decisions about whether to get involved in an aggressive interaction. For example, baboons (*Papio hamadryas ursinus*) were more likely to move towards playback of a grunt call given to recruit support in an aggressive interaction if they had groomed with the caller earlier (mean: 22 min before; range 10–55 min; Cheney et al., 2010). Similarly, vervet monkeys (*Chlorocebus pygerythrus*) were more likely to offer coalitionary support to a groupmate in a conflict if they had groomed together within the last hour (Borgeaud and Bshary, 2015). Other studies have shown that individuals can use knowledge of previous agonistic interactions to inform how best to respond in subsequent aggressive encounters. For instance, chimpanzees (*Pan troglodytes*) that had been involved in an unreconciled conflict earlier in the day (ca. 2 h before) reacted aversively to playback of an aggressive bark from their former opponent’s bond partner (a third-party individual likely to offer aggressive support to the former opponent; Wittig et al., 2014). Moreover, it was recently shown that bystander wasps (*Polistes fuscatus*) were more aggressive towards individuals that they had observed to be less aggressive in a previous (10–30 min earlier) fight with a third party (Tibbetts et al., 2020). It is thus plausible that post-conflict decisions about the avoidance of protagonists and affiliation with groupmates could also occur some time after the relevant contests.

To make behavioural decisions, animals obtain information about social interactions using a variety of sensory modalities. Most research considering social monitoring of within-group conflict has focused on situations where individuals have seen the interaction; hence, bystanders are commonly defined as individuals who have observed the encounter (Schino and Sciarretta, 2015). But for those species living in visually occluded environments, those where group members can be scattered over large distances or those that forage in a way that prevents simultaneous vigilance, acoustic cues can be a valuable source of social information (Bradbury and Vehrencamp, 2011). Numerous species vocalise during or at the end of within-group contests (Bertram et al., 2010; Slocombe et al., 2010). For example, chimpanzees and rhesus monkeys (*Macaca mulatta*) produce screams when experiencing aggression (Gouzoules et al., 1984; Slocombe et al., 2010), whilst little blue penguins (*Eudyptula minor*) give specific calls after a contest is finished (Waas, 1990). These vocalisations likely provide bystanders with valuable information about within-group conflict (Gouzoules et al., 1984; Slocombe and Zuberbühler, 2007; Szipl et al., 2017; Whitehouse and Meunier, 2020). Moreover, they can be used in playbacks to test post-conflict behaviour experimentally.

Here, we investigate experimentally the possibility of delayed post-conflict management decisions in wild dwarf mongooses (*Helogale parvula*); the study population has been habituated to close human presence, facilitating detailed observations and field-based manipulations (Kern and Radford, 2018; Morris-Drake et al., 2019). Dwarf mongooses live in cooperatively breeding groups of up to 30 individuals, comprising a dominant breeding pair (hereafter ‘dominant’ individuals) and non-breeding subordinate helpers (hereafter ‘subordinate’ individuals) of both sexes (Rasa, 1977). Within-group aggressive interactions take two main forms: relatively rare targeted aggression, which usually acts to reinforce rank and is mainly due to reproductive conflict (Rasa, 1977); and relatively common foraging displacements, when a higher-ranking individual displaces a lower-ranking group member from a foraging patch and steals their prey (Sharpe et al., 2016, 2013). Foraging displacements generally involve the following behavioural sequence: the higher-ranking individual produces deep growls as it approaches the lower-ranking group member; the former then hip-slams the latter away from the food resource; and the displaced individual typically produces high-pitched squeals whilst it retreats (Sharpe et al., 2016, 2013). Previous work has shown that dwarf mongooses can use vocal information to facilitate delayed rewarding of cooperative contributions by groupmates (Kern and Radford, 2018). We now test whether vocal cues of within-group conflict elicit delayed behavioural responses (avoidance or changes in affiliation) by non-participant group members.

## Results

We initially used both observational data and a playback experiment to determine whether there is evidence that bystanders take notice of conflict between groupmates and if they engage in affiliative interactions (grooming or vocal exchanges) in the immediate aftermath (full details in *Materials and Methods*). To collect data relating to natural foraging displacements (which occur at a mean±SE observer-detected rate of 2.6±0.2 events per 3-h observation session, range=0–10, N=127 observation sessions across eight groups), we conducted focal watches on foraging subordinates in two situations: immediately after the human observer heard a foraging displacement (conflict situation) and on a matched occasion when there had been no foraging displacement for at least 10 min (control situation). Paired data were collected from 16 subordinates in six groups, with conflict and control focal watches counterbalanced in order between individuals. To test experimentally the immediate responses of bystanders, and to isolate the importance of foraging-displacement vocalisations as a cue to conflict occurrence, we presented 17 foraging subordinates in eight groups with two playback treatments in a matched, counterbalanced design (Experiment 1). The conflict treatment entailed initial playback of close calls from a dominant individual and a subordinate individual from the same group as the focal individual, followed by playback of the dominant growling and the subordinate squealing (simulating a foraging displacement); the control treatment entailed the playback of close calls from the same two individuals for the same duration as a full conflict-treatment playback track (Fig. 1). We chose for playback the combination of a dominant individual as the aggressor and a subordinate individual as a victim because this is the most common dyadic pairing observed in natural foraging displacements (74.3% of 740 events in 12 groups).

**Fig. 1.**
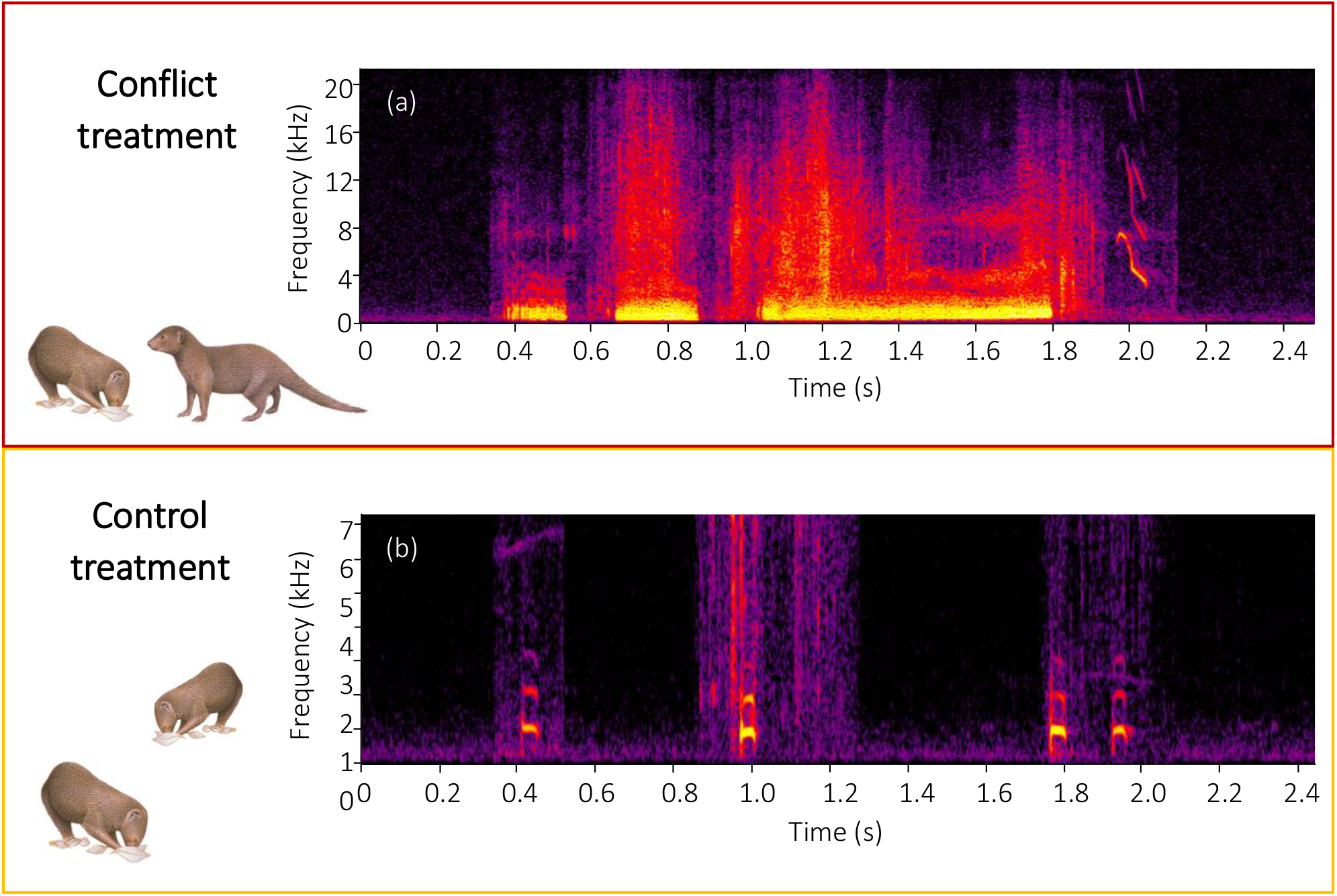
Spectrograms of the final sections of example (a) conflict and (b) control playback tracks. A conflict track concluded with three growls from a dominant aggressor followed by a squeal from a subordinate victim, whilst a control track concluded with three close calls from the same dominant individual followed by one close call from the same subordinate individual as in the matched conflict track. Spectrograms were created in Raven Pro 1.5 using a 1024 point fast Fourier Transform (Hamming window, 75% overlap, 2.70 ms time resolution, 43 Hz frequency resolution).

We found evidence that bystanders take notice of conflicts between groupmates but no indication of immediate post-conflict affiliative exchanges with either the protagonists or other group members. In the 2–3 min following both natural foraging displacements (Wilcoxon signed-rank test: Z=3.154, N=16, Monte Carlo P<0.001; Fig. 2a) and those simulated by playback (Z=3.527, N=17, P<0.001; Fig. 2b), focal foragers spent a significantly greater proportion of time vigilant than in matched-control, non-conflict situations. This was because individuals were conducting vigilance bouts both at a significantly greater rate (observational data: Z=2.517, N=16, P=0.008; experimental data: Z=3.479, N=17, P<0.001) and for significantly longer durations (observational: Z=2.500, N=15, P=0.009; experimental: Z=3.574, N=17, P<0.001) in the aftermath of a foraging displacement compared to control periods. However, the focal individual did not engage in any post-conflict grooming in the 5 min following either natural or simulated foraging displacements; grooming is generally rare during foraging periods in dwarf mongooses (Kern and Radford, 2018). There was also no evidence of vocal ‘grooming-at-a-distance’ (Arlet et al., 2015; Kulahci et al., 2015), as the close-call rate of focal individuals was not significantly greater following natural (Z=1.500, N=16, P=0.144) or simulated (Z=1.491, N=17, P=0.144) foraging displacements compared to control situations. The increased vigilance following foraging displacements indicates that other group members have noticed their occurrence; the experimental results demonstrate that the vocal cues are sufficient to trigger this reaction. However, there is no evidence that dwarf mongoose bystanders engage in post-conflict affiliative behaviour in the immediate aftermath of hearing a within-group contest.

**Fig. 2.**
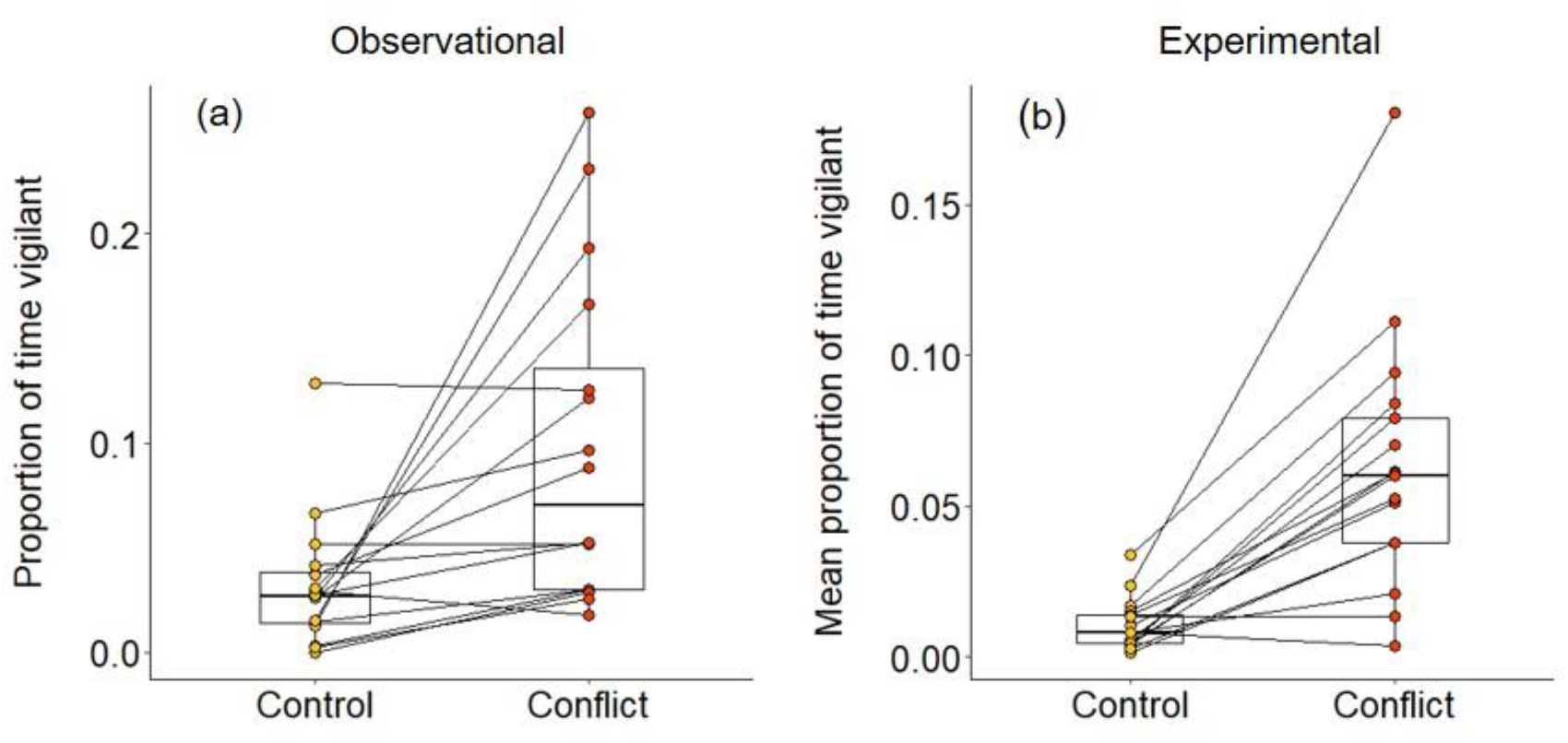
Immediate effect of within-group conflict on dwarf mongoose vigilance behaviour. Compared to control situations, (a) natural foraging displacements (observational; N=16 individuals in six groups) and (b) simulated foraging displacements (experimental; N=17 individuals in eight groups) both resulted in a greater proportion of time spent vigilant by foragers in the subsequent 2–3 min. Shown in both panels are boxplots with the median and quartiles; whiskers represent data within quartiles ± 1.5 times the interquartile range. Values for each individual are given as circles, with lines connecting data from the same individual; in some instances, more than one individual has the same value, hence the number of lines can appear less than the stated sample size.

To test if there were delayed effects of within-group conflict on affiliative behaviour (grooming), we conducted a second repeated-measures playback experiment on eight groups (Experiment 2, Fig. 3; full details in *Material and Methods*). The general experimental design followed (Kern and Radford, 2018). In each trial session, we either simulated: an increase in the conflict between a dominant (aggressor) and a subordinate (victim) group member through playback of their foraging-displacement vocalisations (conflict treatment); or played back just the close calls of those individuals for an equivalent period (control treatment). Trials were on separate days with treatment order counterbalanced between groups. In each trial, 6–9 playbacks (mean±SE: 8.5±0.2, N=16 trials) were carried out over the course of 3 h in the afternoon whilst the group were foraging and before they moved towards their evening sleeping refuge (mean±SE period between final playback and first grooming bout at the sleeping refuge: 37±5 min, N=16 trials); individual playbacks were as in Experiment 1 with different tracks played each time. At the refuge, we collected data *ad libitum* on all adult grooming interactions, including the identity of those involved and bout duration; each bout was always between just two individuals and generally mutual (both parties approaching each other and grooming, without an obvious initiator). If within-group conflict does have delayed effects on affiliative behaviour, we expected an increase in the occurrence of foraging displacements to result in changes in evening grooming levels; 90% of grooming bouts occur at the sleeping refuge (N=6,376 bouts, 174 individuals; Kern and Radford, 2018).

**Fig. 3.**
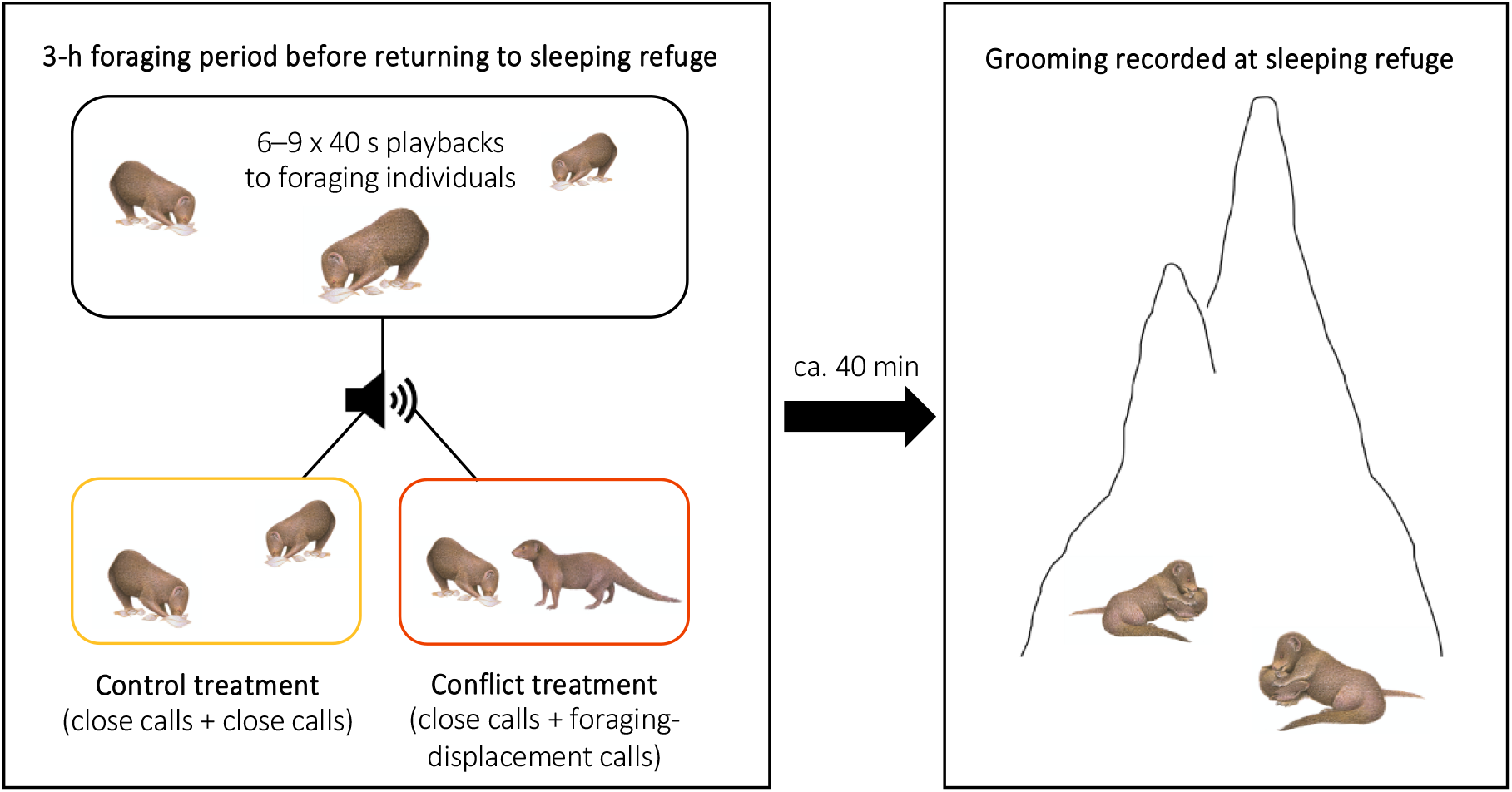
Illustration of the protocol for Experiment 2. Within-group conflicts between a dominant aggressor and a subordinate victim were simulated during conflict-treatment afternoons using playback of foraging-displacement calls, with only close calls of the same individuals played back in control sessions. All grooming at the evening sleeping refuge was subsequently recorded following both treatments.

Overall, we found that group members were significantly less likely to be involved in grooming interactions in the evenings following conflict afternoons compared to control afternoons (generalised linear mixed model [GLMM]: χ^2^=5.401, df=1, P=0.020; Table 1a; Fig. 4a). However, when considering only those individuals that engaged in grooming, they spent a significantly greater proportion of time doing so on evenings when there had been an earlier simulated increase in conflict compared to control evenings (linear mixed model [LMM]: χ^2^=16.522, df=1, P<0.001; Table 1b; Fig. 4b). This was because these individuals were grooming more frequently (χ^2^=14.810, df=1, P<0.001; Table 1c) and for longer per bout (χ^2^=3.958, df=1, P=0.047; Table 1d) after a simulated increase in conflict compared to control conditions. These results indicate that there is an overall response to simulated conflict within the group, but we also made some specific predictions. Assuming that aggressors and victims can be identified from their vocalisations—which has been demonstrated for dwarf mongoose close calls (Sharpe et al., 2013), recruitment calls (Kern and Radford, 2016) and surveillance calls (Kern and Radford, 2018)—we predicted that subordinates might engage in either less grooming (due to wariness) or more grooming (as possible appeasement) with aggressors, and that they might engage in more grooming with victims (as possible consolation).

**Fig. 4.**
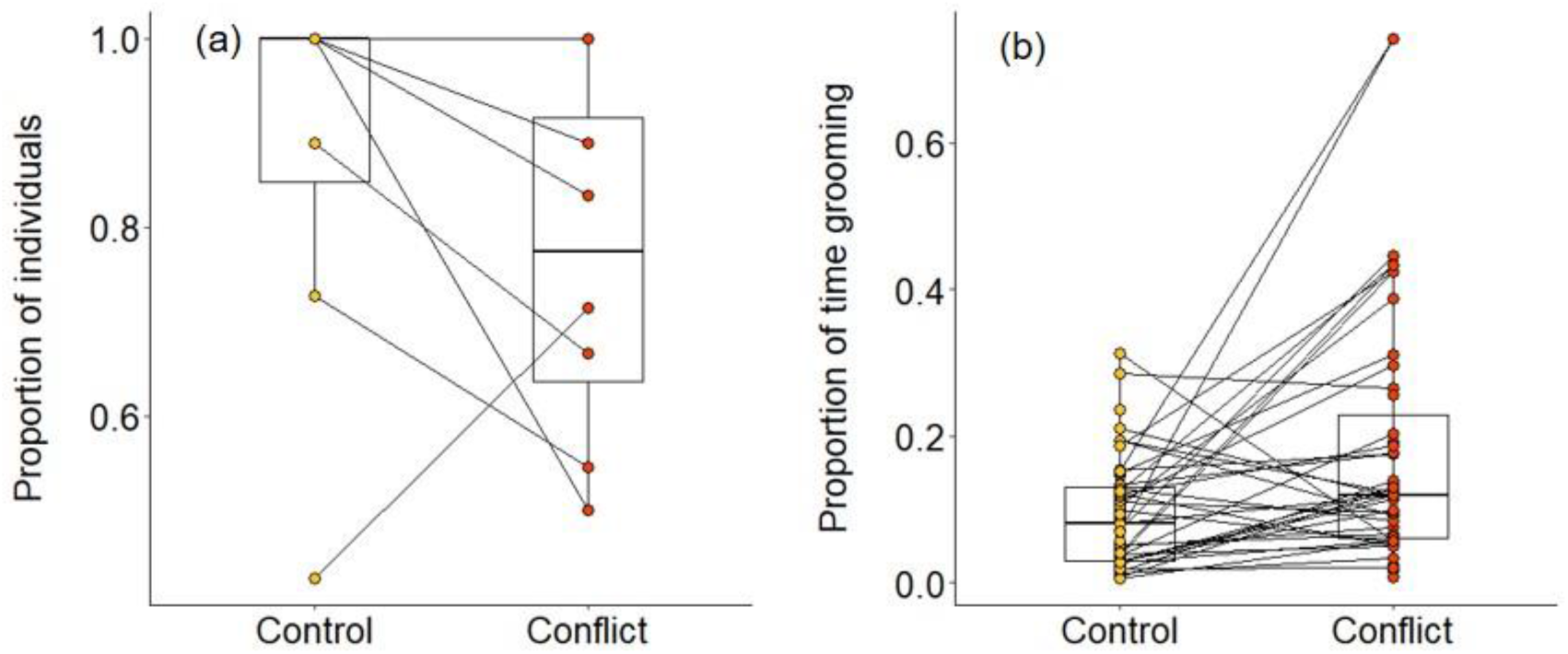
Delayed effect of experimentally increased within-group conflict on dwarf mongoose grooming behaviour. Compared to control afternoons, those with simulated additional foraging displacements resulted in (a) a smaller proportion of adult group members engaged in evening grooming behaviour (N=8 groups) but (b) a greater proportion of time engaged in grooming by those individuals that did any grooming (N=63 individuals in eight groups). Shown in both panels are boxplots with the median and quartiles; whiskers represent data within quartiles ± 1.5 times the interquartile range. Values for each group or individual are given as circles, with lines connecting data from the same group or individual; orphan points, where an individual only groomed in one treatment, are also plotted. In some instances, more than one group or individual has the same value, hence the number of lines can appear less than the stated sample size.

**Table 1.** Output from (a) a GLMM and (b–d) LMMs investigating the grooming behaviour of adult dwarf mongooses at the evening refuge. All models contained treatment (conflict, control) as a fixed effect (the reference level in the table is ‘conflict’), with Individual ID nested within Group ID as random effects. The GLMM (binomial error distribution and logit-link function) examined whether an individual was involved in a grooming bout (Yes or No). Subsequent LMMs focused on those individuals that did participate in grooming, examining (b) the log-transformed proportion of time spent grooming, (c) the log-transformed rate of grooming interactions and (d) the log-transformed mean grooming-bout duration. Significant fixed effects shown in bold; variance ±SD reported for random effects (in italics).

| | Effects | Estimate±SE | df | $\chi^2$ | P |
| --- | --- | --- | --- | --- | --- |
| **(a) Individual involvement in grooming** |  |  |  |  |  |
| Random effects | *Group ID* | 0.919±0.959 | | | |
| | *Individual ID in Group* | <0.001 ± <0.001 | | | |
| Minimal model | (Intercept) | 1.267±0.497 | | | |
|  | **Treatment (Conflict)** | **1.106±0.488** | **1** | **5.401** | **0.020** |
| **(b) Proportion of time spent grooming** |  |  |  |  |  |
| Random effects | *Group ID* | 0.130±0.360 | | | |
| | *Individual ID in Group* | 0.308±0.555 | | | |
| Minimal model | (Intercept) | -2.114±0.186 | | | |
|  | **Treatment (Conflict)** | **-0.681±0.155** | **1** | **16.522** | **<0.001** |
| **(c) Rate of grooming bouts** |  |  |  |  |  |
| Random effects | *Group ID* | 0.069±0.262 | | | |
| | *Individual ID in Group* | 0.124±0.353 | | | |
| Minimal model | (Intercept) | -1.269±0.139 | | | |
|  | **Treatment (Conflict)** | **-0.515±0.125** | **1** | **14.810** | **<0.001** |
| **(d) Mean grooming-bout duration** |  |  |  |  |  |
| Random effects | *Group ID* | 0.047±0.217 | | | |
| | *Individual ID in Group* | 0.067±0.258 | | | |
| Minimal model | (Intercept) | 3.257±0.104 | | | |
|  | **Treatment (Conflict)** | **-0.167±0.083** | **1** | **3.958** | **0.047** |

We found strong evidence that simulating aggressive behaviour by a dominant individual during the afternoon resulted in subordinates engaging in less grooming with it at the sleeping refuge that evening. Following conflict trials, subordinates groomed with the dominant pair for a smaller proportion of time than after control trials (Wilcoxon signed-rank test: Z=2.240, N=8, P=0.021). The reduced affiliative engagement by subordinates was driven by a change in behaviour towards the simulated aggressor specifically: there was significantly less grooming between subordinates and the simulated aggressor on conflict evenings compared to control evenings (Z=2.521, N=8, P=0.008; Fig. 5a), but no such treatment difference in the proportion of time that subordinates groomed with the dominant whose calls were not played back (Z=0.105, N=8, P=1; Fig. 5b). At least in part, this difference was due to there being a significantly smaller proportion of subordinates grooming with the simulated aggressor in the evening of conflict trials compared to control trials (Z=2.201, N=8, P=0.033; Fig. 5c), but no such treatment difference in the proportion of subordinates that groomed with the non-playback dominant (Z=0.813, N=8, P=0.499; Fig. 5d). Moreover, on those occasions where individuals did groom, bout durations were somewhat shorter on conflict evenings compared to control evenings for grooming involving simulated aggressors (mean±SE duration, post-control: 34±11 s; post-conflict: 23±5 s; N=4 pairs of trials), while the reverse was true for grooming involving the matched dominant (post-control: 28±8 s; post-conflict: 34±8 s; N=4 pairs of trials); small sample sizes precluded statistical analysis. To our knowledge, this is the first evidence for a reduction in grooming of aggressors by bystanders; some previous studies have documented increased grooming of aggressors by bystanders in the immediate aftermath of a single contest (Cordoni and Palagi, 2015; Palagi et al., 2008; Pallante et al., 2018), whilst a few others have found no evidence for such an increase (Judge, 1991; Romero et al., 2008; Verbeek and de Waal, 1997). Subordinate bystanders could be avoiding the aggressor to reduce the likelihood of redirected aggression, which parallels the main strategy employed in the immediate aftermath of contests by meerkat (*Suricata suricatta*) and rook (*Corvus frugilegus*) victims attempting to avoid renewed aggression (Benkada et al., 2020; Kutsukake and Clutton-Brock, 2008).

**Fig. 5.**
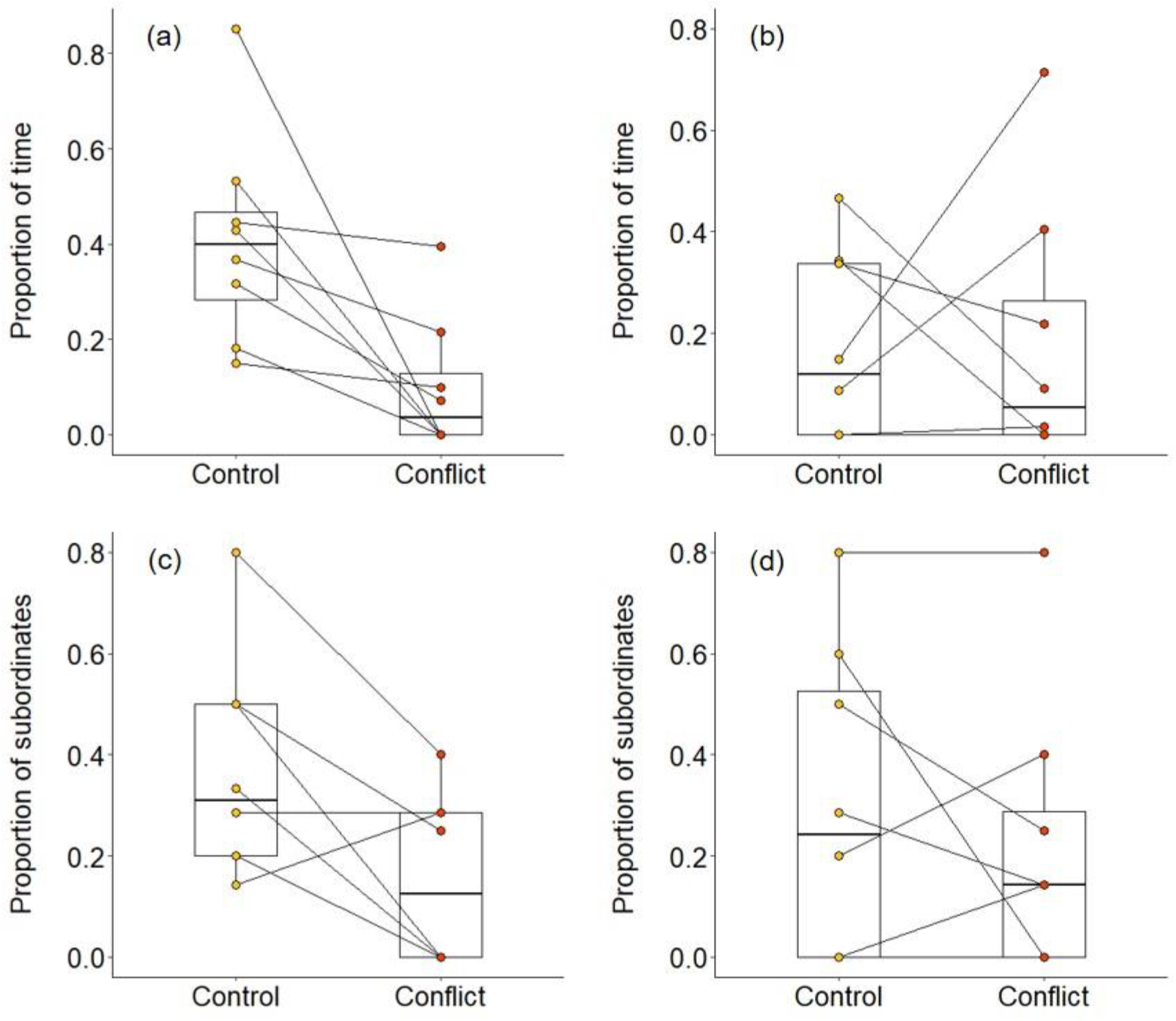
Delayed effect of experimentally increased within-group conflict on the grooming between subordinate bystanders and the simulated aggressor. Compared to control afternoons, those with simulated additional foraging displacements between a dominant aggressor and a subordinate victim resulted in (a) a smaller proportion of time engaged in evening grooming by subordinate bystanders with the dominant aggressor, but (b) no such treatment difference in the proportion of time that subordinate bystanders groomed with a non-playback dominant. At least in part, this was because (c) a smaller proportion of subordinate bystanders groomed with the dominant aggressor in the evening of conflict afternoons compared with control ones, but (d) there was no such treatment difference in the proportion of subordinates involved in grooming with a non-playback dominant. Shown in all panels are boxplots with the median and quartiles; whiskers represent data within quartiles ± 1.5 times the interquartile range. Values for each group are plotted separately (N=8), with lines connecting data from the same group; in some instances, more than one group has the same value, hence the number of lines can appear less than eight.

We also found some evidence that increasing within-group conflict during the afternoon resulted in more evening grooming between subordinates. When considering all bouts between subordinate group members, there was no significant treatment difference in the proportion of time spent grooming (Wilcoxon signed-rank test: Z=1.540, N=8, P=0.146), but subordinate–subordinate grooming bouts were, on average, significantly longer on conflict evenings compared to control evenings (Z=2.366, N=7, P=0.015; Fig. 6a). Considering bouts involving particular individuals, there were indications that victims might receive a conflict-driven increase in grooming from other subordinates not seen for preselected control subordinates (those whose squeals had not been played back), but no statistically significant differences. The proportion of time grooming that involved the simulated victim was doubled on conflict evenings (mean±SE: 0.31±0.09) compared to control evenings (0.15±0.06; Z=1.572, N=8, P=0.156; Fig. 6b), whereas there was, if anything, a decrease for the preselected control subordinate (control: 0.37±0.12; conflict: 0.28±0.09; Z=0.280, N=8, P=0.843; Fig. 6c). The treatment difference in mean bout duration was also greater for grooming involving simulated victims (36±14 s, N=3 pairs of trials) than that involving preselected control subordinates (22±24 s, N=3 pairs of trials), but too few matched evenings involved the relevant individuals to allow statistical testing. The increase in the average duration of subordinate–subordinate grooming is in-line with the increase in bystander–bystander grooming seen in some species in the immediate aftermath of a contest (Judge and Mullen, 2005). Such affiliation could reduce the group-wide social anxiety induced by aggression (De Marco et al., 2010; Judge and Bachmann, 2013; Schino and Sciarretta, 2015).

**Fig. 6.**
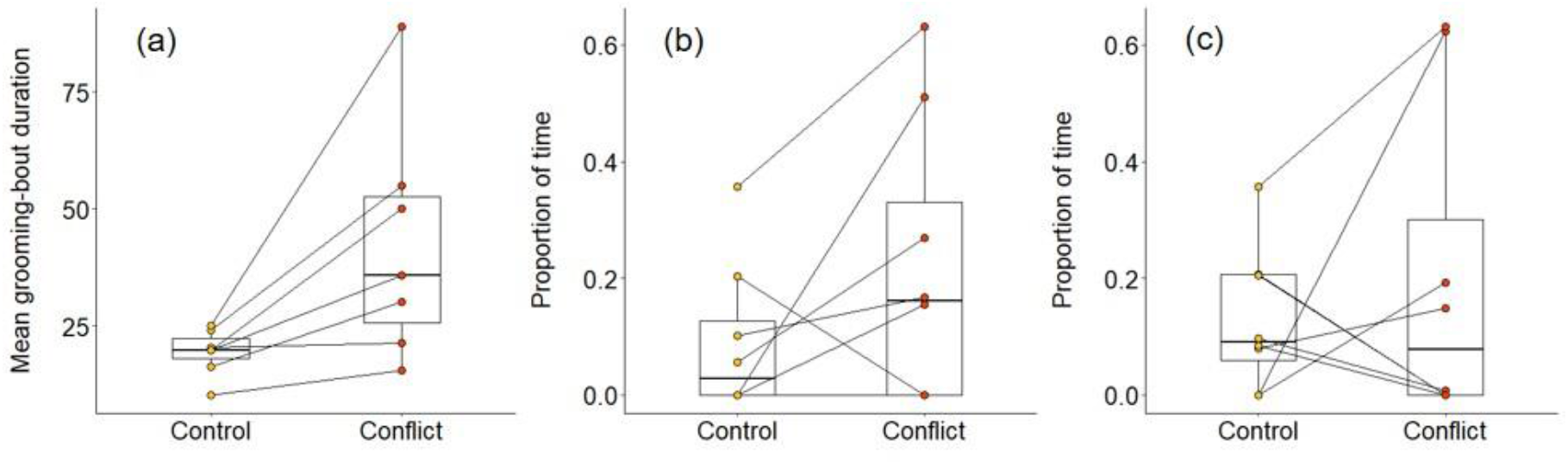
Delayed effect of experimentally increased within-group conflict on the grooming between subordinate bystanders. Compared to control afternoons, those with simulated additional foraging displacements between a dominant aggressor and a subordinate victim resulted in (a) a greater mean duration of grooming bouts (s) between subordinate group members. There was (b) an indication that subordinate bystanders and victims groomed for a greater proportion of time on conflict evenings compared to control ones, although the result was not statistically significant; (c) there was no equivalent treatment difference in the proportion of time that subordinates groomed with non-victim subordinates. Shown in all panels are boxplots with the median and quartiles; whiskers represent data within quartiles ± 1.5 times the interquartile range. Values for each group are plotted separately (N=8), with lines connecting data from the same group; in some instances, more than one group has the same value, hence the number of dashed lines can appear less than eight.

## Discussion

We believe that we provide compelling experimental evidence for delayed post-conflict management behaviour. Dwarf mongoose bystanders did not engage in any obvious post-conflict affiliation in the immediate aftermath of natural or simulated foraging displacements involving a dominant and subordinate group member, but did adjust their later grooming behaviour at the evening sleeping refuge following a simulated increase in within-group conflict during the afternoon. Numerous studies on a range of species have found changes in affiliative behaviour between various combinations of protagonists and bystanders in the minutes after within-group contests (Aureli et al., 2002; de Waal, 2000). It remains unknown whether, in those species, there might be delayed effects (as we have found) of those contests that are not resolved immediately. By using call playbacks, we likely did not alter the state or behaviour of the individuals who were simulated to be the aggressor and victim. Consequently, we can rule out the possibility that differences in grooming result from experimentally induced satiation effects (which might have been the case if we had caused foraging displacements with the presentation of food items; Sharpe et al., 2013). Moreover, the use of playback simulations, rather than generation of actual contests, means that the delayed grooming effects are most likely driven by subordinate bystanders behaving differently towards perceived protagonists, rather than solicitation or rejection of grooming by the latter. Overall, our results demonstrate that individuals can retain information relating to earlier within-group conflict and use it when making later decisions about management-related behaviours.

Our experiments show that dwarf mongooses can extract information about within-group conflict, and the identity of at least some protagonists, from vocal cues alone. This adds to a growing body of work demonstrating the ability of social species to garner information acoustically about aggressive interactions (Gouzoules et al., 1984; Slocombe et al., 2010; Slocombe and Zuberbühler, 2007); for example, male little blue penguins had an increased heartrate after hearing vocalisations produced by winners of a contest compared to those produced by losers (Mouterde et al., 2012). Our findings also complement the small number of studies showing that social animals use vocalisations to assess the behaviour, such as the reliability (Blumstein et al., 2004) and cooperative contributions (Kern and Radford, 2018), of individually identifiable groupmates. Acoustic monitoring is beneficial as it allows information acquisition in environments where it would be difficult to do so visually (e.g. in low-light and dense vegetation) or when group members are widely scattered and communication is needed over long distances (Bradbury and Vehrencamp, 2011). Moreover, acoustic information can be gathered at a relatively low cost: it can be done whilst still actively foraging (Hollén et al., 2008) and, in the case of aggressive encounters, at a safe distance that minimises the risk of the information-gatherer receiving any redirected aggression. Monitoring behaviours acoustically is likely not possible for all within-group interactions (e.g. grooming) or in all social systems, but the calls commonly produced during and at the end of aggressive contests (Bertram et al., 2010; Slocombe et al., 2010) provide a valuable means for bystanders to inform subsequent decision-making.

We found strong evidence that subordinate bystanders engage in less grooming with simulated aggressors, but whether they increased their grooming with the simulated victim was less clear-cut. There are several possible explanations for this difference in the strength of response exhibited to the two protagonists. First, all subordinates might be wary of the aggressor and so potentially reduce their grooming with that individual, whereas perhaps only those who are strongly bonded to the victim might engage in extra grooming with it (Fraser et al., 2009, 2008); any such victim-related effect might be diluted by considering all subordinates in analyses. Strong within-group relationships are apparent in dwarf mongoose groups (Kern and Radford, 2016), but we do not have the power in this study to consider how relationship quality influences delayed post-conflict grooming. Another possible reason for the difference in grooming responses to aggressors and victims is that there could be selective attention towards high-ranking individuals (Chance, 1967). Many primate species, for example, focus attention on higher-ranking groupmates or those with whom they have an antagonistic relationship (Keverne et al., 1978; McNelis and Boatright-Horowitz, 1998), possibly to avoid aggression (Schino and Sciarretta, 2016). Since our simulated aggressors were dominants and our simulated victims were subordinates, the stronger effect of increased conflict on grooming with the former could reflect such an attention bias. Alternatively, our results could be driven by differences in the natural acoustic properties of aggressive growls and submissive squeals (Gustison and Townsend, 2015). In principle, squeals might encode less identity information than growls (Owren and Rendall, 2003; Rendall et al., 1996), although a number of studies have found that calls similar in structure and function to dwarf mongoose squeals are individually identifiable (Cheney and Seyfarth, 1980; Fischer, 2004; Gouzoules et al., 1984; Slocombe and Zuberbühler, 2005). In addition, our playback contained three growls and one squeal (to reflect natural foraging displacements), which could have made growls more salient or memorable and/or aided easier discrimination of the aggressor compared to the victim. It might also be more cognitively demanding for receivers to discriminate the squeals from multiple subordinate individuals in a group, compared to growls, which are highly likely to come from one of the two dominant individuals. Finally, since contest-related vocalisations may vary depending on the severity of an attack (Gouzoules et al., 1984; Slocombe and Zuberbühler, 2007), it is possible that we used less salient squeals than growls in our playbacks (although both were recorded during natural foraging displacements). Future work is required to tease these possibilities apart.

In summary, our results demonstrate that dwarf mongooses can obtain information about within-group contests (including protagonist identity) acoustically, retain that information and use it to inform decisions about conflict management with a temporal delay. Such a delay might be most apparent in situations where there is little opportunity for immediate post-contest affiliation (as is the case with foraging dwarf mongooses); it may also be most apparent when there is a cumulative build-up of unresolved conflict. There is increasing experimental evidence that social animals can remember past events and take these into account when deciding whether to get involved in a contest (Borgeaud and Bshary, 2015; Cheney et al., 2010; Tibbetts et al., 2020; Wittig et al., 2014); we demonstrate that this ability extends to post-conflict affiliative behaviour. The cognitive demands of tracking individuals and their behaviours, remembering that information and using it when making decisions is why social interactions within (Dunbar and Shultz, 2007) and between(Ashton et al., 2020) groups are believed to be strong drivers of animal intelligence.

## Materials and Methods

### Study site and population

We conducted our study on Sorabi Rock Lodge (24° 11’S, 30° 46’E), a private game reserve in the Limpopo Province, South Africa; full details in Kern and Radford, 2013.This is the site of the Dwarf Mongoose Research Project (DMRP), which has been studying a wild population of dwarf mongooses since 2011. At the time of study (June to October 2019; non-breeding season), eight dwarf mongoose groups (mean±SE group size: 12.3±1.7, range: 5–16) were fully habituated to the close presence (<5 m) of human observers on foot. All the individuals in the population were identifiable, either through dye-marks on their fur (blond hair dye applied using an elongated paintbrush) or natural features, such as scars. Individuals older than 1 year were classified as adults (Kern et al., 2016); data collection focused on adults as younger individuals are seldom involved in foraging displacements. Adults were sexed by observing ano-genital grooming (Kern et al., 2016) and classified as being either dominant (the male and female breeding pair) or subordinate; dominance status was established through observation of targeted aggression, scent marking and reproductive behaviour (Kern and Radford, 2013; Rasa, 1977).

### Observational data collection

To determine the natural frequency of foraging displacements in our experimental period, we recorded all observer-detected occurrences of such behaviour during observation sessions; this included displacements that were seen and heard. The calculated rate is likely a conservative estimate as an observer could have missed a foraging displacement (particularly when the group was relatively widely scattered). We used data collected *ad libitum* as part of the long-term DMRP to assess the likelihood of particular dyads of individuals (aggressor–victim: dominant–dominant, dominant–subordinate, subordinate–subordinate, subordinate–dominant) being involved in a foraging displacement.

To collect data on responses to natural foraging displacements, we conducted paired focal watches (conflict and control) of 2–3 min duration on 16 subordinate group members in six groups whilst they were foraging; conflict and control focal watches did not differ significantly in their duration (Wilcoxon signed-rank test: Z=0.952, N=16, P=0.380). A conflict watch was carried out immediately after a foraging displacement was heard by the observer, whilst a control watch was carried out when there had been no foraging displacement (or any other agonistic interaction) for at least 10 min. We only carried out focal watches when the relevant mongoose was in medium-cover habitat (20-60% ground cover), weather conditions were calm (still or light breeze), there had been no alarm call (conspecific or heterospecific in the previous 10 min, there had been no predator encounter or inter-group interaction for at least 30 min, and the focal individual was not on the periphery of the group. We abandoned focal watches, and repeated them later, if the focal individual stopped general foraging activities or if there was an alarm call within the first 2 min. Otherwise, we aimed to collect 3 min of uninterrupted data, but if a behavioural change or alarm call occurred between the second and third minute, then the focal watch was retained. Pairs of watches on the same focal individual were completed within 1 month (mean±SE: 8.1±2.7 days apart, range: 0–30 days); group composition always remained the same between a pair of watches, and a minimum of 1 h was left between watches that were conducted on the same day. We watched nine individuals first in control conditions and seven first following a foraging displacement.

During each focal watch, we recorded behavioural data to a Dictaphone (ICD-PX312, Sony, Sony Europe Limited, Surrey, UK). Dwarf mongooses have two types of vigilance behaviour: vigilance scans, where individuals temporarily stop foraging in a head-down position to scan their surroundings (Kern et al., 2016); and sentinel behaviour, where individuals cease foraging to scan from a raised position (minimum 10 cm above ground level; Kern and Radford, 2013). They also produce low-amplitude close calls continuously whilst foraging (Kern and Radford, 2013; Sharpe et al., 2013). Throughout each focal watch, we dictated the start and end point of each vigilance scan and sentinel bout, along with the occurrence of each close call and any grooming interaction with a groupmate. These data were used to calculate the proportion of time spent vigilant, the vigilance rate, the mean duration of vigilance bouts and the close-call rate; no grooming occurred during these focal watches. No individuals acted as a sentinel during the observational focal watches and therefore the above vigilance response measures were based on scan data only. We used Wilcoxon signed-ranks tests to analyse the dependent variables in SPSS 24 (IBM Corp, 2016); due to small sample sizes, we used the Monte Carlo resampling method (based on 10,000 samples) to generate P-values.

### Experimental stimuli

We conducted two field-based repeated-measures experiments using playbacks to simulate the occurrence of conflict between group members. Each experiment involved the playback of ‘conflict’ and ‘control’ tracks. We recorded all calls for track creation when weather conditions were calm using a Marantz PMD660 professional solid-state recorder (Marantz America, Mahwah, NJ) connected to a handheld Sennheiser ME66 directional microphone (Sennheiser UK, High Wycombe, Buckinghamshire, UK) with a Rycote softie windshield (Rycote Microphone Windshields, Stroud, Gloucestershire, UK). The Marantz was set to record at 48 kHz with a 16-bit resolution, and files were saved in wav format. For conflict tracks, we recorded aggressive growls and submissive squeals opportunistically from natural foraging displacements or from conflicts induced by the presentation of a small amount of hard-boiled egg. Growls were recorded from either the dominant male or dominant female in each group and squeals were recorded from a subordinate male or female in each group; all recorded calls came from foraging displacements where the dominant was the aggressor and the subordinate was the victim. We recorded close calls, for use in both control and conflict tracks, from the same dominant and subordinate individuals whilst they were foraging. Recordings of all vocalisations were made 0.5–5 m from the relevant individual.

We formed 40 s playback tracks in Audacity (version 2.1.3) by extracting calls of good signal-to-noise ratio from original recordings and inserting them into ambient-sound recordings; ambient sound was recorded from the centre of the territory of the focal group on calm days and in the absence of dwarf mongooses. The first 36 s of each track (conflict and control) consisted of non-overlapping close calls from the relevant dominant and subordinate individual, with a rate of 1 close call every 6 seconds per individual. This rate of close calling falls within the natural range (Kern and Radford, 2013). For conflict tracks, the last 4 s consisted of a sequence of three growls from the dominant followed by one squeal from the subordinate; multiple growls and a single squeal reflects natural foraging displacements (personal observation). In control tracks, the last 4 s consisted of three close calls from the dominant followed by one close call from the subordinate, to match the number of vocalisations in conflict tracks. Individual tracks always contained vocalisations from same-sex individuals.

We created nine unique conflict and control tracks for each group. Given that the first 36 s of each track comprised close calls from the dominant and subordinate individual, we created three close-call sequences for each individual (each sequence contained six close calls), resulting in nine unique close-call combinations. For the conflict tracks, in which the last 4 s contained growls and a squeal, we created three different growl sequences for the dominant (each sequence consisted of three growls), which were each combined with three separate squeals from the subordinate. Lastly, for the final 4 s of the control tracks, we made three close-call sequences for the dominant (each sequence contained three close calls to match the number of growls in conflict tracks), and combined these with three different close calls from the subordinate. We applied a low-pass filter (set to 200 Hz) to all tracks to remove low-frequency disturbance.

We played back tracks from an iPhone (Apple, Cupertino, CA), connected to a Rokono B10 (London, UK) portable loudspeaker concealed in vegetation. We set the amplitude to a sound-pressure level of 55 dB(A) at 1 m for close calls and growls, and 65 dB(A) at 1 m for squeals. This was the relevant amplitude of these vocalisations as determined by measurement of natural calls with a HandyMAN TEK 1345 sound-level meter (Metrel UK Ltd., Normanton, UK).

### Experiment 1 protocol

Experiment 1 was a complement to the observational focal watches (see *Observational data collection*), aiming to test whether bystanders might garner information about within-group conflict solely from vocalisations. We randomly selected 17 subordinate individuals (excluding those whose calls were used in the playback tracks) to receive the two treatments (conflict and control) on separate days and in a counterbalanced order. Each treatment was repeated 2–3 times per individual during the same observation session, using a different playback track each time, with a minimum of 10 min between repeats; for one individual, it was possible to run one of the treatments only once. We completed the two treatments for the same individual within 2 weeks of each other (mean±SE: 2.8±0.7 days apart, range: 1–11 days) and at the same time of day (either between 07:00 and 12:00 or between 12:30 and 17:30). The 17 focal individuals were from eight groups; for groups where there was more than one focal individual (N=4 groups), we completed both treatments on one individual before moving on to the next.

We conducted playbacks when the focal individual was foraging in medium habitat with little or no breeze and when the callers in the playback were not the focal individual’s nearest neighbour (other pre-requisites detailed in *Observational data collection*). Where possible, we placed the loudspeaker in the general direction of the playback individuals. As soon as the playback finished, we conducted a 2–3 min focal watch; the mean duration of focal watches was not significantly different between treatments (Wilcoxon signed-rank test: Z=1.397, N=17, P=0.168). Collection of vigilance and close-calling data was identical to that for observational focal watches.

We analysed the same response variables as those for the natural foraging displacements (see *Observational data collection*): proportion of time spent vigilant, vigilance rate, mean duration of vigilance bouts and the close-call rate; no grooming occurred in any focal watches. Since each treatment was repeated 2–3 times on an individual, we analysed the mean for each response measure; Wilcoxon signed-ranks tests were used. In five out of 94 trials, an individual acted as a sentinel. We therefore ran the vigilance response measures including and excluding this sentinel behaviour. The data reported in the *Results* section are those excluding sentinel bouts, but qualitatively similar results were found for those including this behaviour.

### Experiment 2 protocol

Experiment 2 aimed to test whether there was a delayed effect of within-group conflict on affiliation between group members. We gave eight groups two treatments each on separate days, with treatment order counterbalanced between the groups. On conflict days, the perceived level of within-group conflict was increased during the afternoon by playback of up to nine conflict tracks. On control days, perceived levels of within-group conflict were unmanipulated; up to nine control tracks were played back during the afternoon instead. There was no treatment difference in the number of natural foraging displacements that occurred throughout the afternoon (Wilcoxon signed-rank test: Z=1.725, N=8, P=0.158). We completed the two treatments to the same group within 2 weeks of each other (mean±SE: 3.3±1.0 days apart, range: 1–9 days). Trials were only attempted when the weather conditions were suitable (not too windy or cold) and were abandoned if any major disturbances occurred during the afternoon (e.g. predation attempts, inter-group interactions, multiple latrine events).

On a trial afternoon, we played back tracks from the centre of the foraging group approximately every 20 min during the 3-h period before the group started moving to an evening sleeping refuge. There were five trials (2 conflict, 3 control) where circumstances (e.g. groups on the move, individuals foraging too far apart) prevented us from completing all nine planned playbacks in an afternoon (mean±SE number of playbacks per trial: 8.5±0.2, range: 6–9) before the group headed to their sleeping refuge. Once at the refuge (always termite mounds), we recorded all instances of adult grooming behaviour *ad libitum* until the mongooses went below ground for the night; it is possible to collect data on all group members simultaneously because they are within a small area around the refuge compared to being scatted more widely when foraging (i.e. in Experiment 1). Data collection involved dictating the identity of grooming partners and the start and end point of each bout. Periods of grooming data collection at the refuge (mean±SE: 15.5±2.3 min, range: 2–37 min) were not significantly different in duration between treatments (Wilcoxon signed-rank test: Z=1.332, N=8, P=0.209).

To analyse the overall grooming data at the refuge (including grooming bouts >5 s; Kern and Radford, 2018), we constructed mixed models in RStudio 3.6.2 (R Core Team 2019) using the package lme4 (Bates et al., 2015). For all models, we included treatment as a fixed effect and nested Individual ID within Group ID as random effects to account for data from the same individuals and groups. Error distributions were chosen such that there were no deviations from normality or homoscedasticity, as checked by graphical examination of residual plots; certain response variables were transformed to meet the assumptions of parametric testing (see below). To assess the significance of treatment (our one fixed effect), we compared a model containing treatment to a model without it (null model) using a likelihood ratio test (ANOVA model comparison, χ^2^ test) and comparison of the Akaike Information Criterion. All tests were two-tailed and considered significant below an alpha level of 0.05.

We first ran a GLMM with a binomial error distribution and logit-link function to assess whether there was a difference in the number of adult individuals that participated in grooming behaviour; our response measure was a binary term – did the individual engage in any grooming (Yes or No). For those individuals that did participate in grooming, we ran Gaussian LMMs to understand this behaviour further. We first analysed the log-transformed proportion of time that individuals spent grooming (summed grooming durations for each individual divided by the time available for grooming at the refuge, with the latter defined as the duration between the first and last grooming bout). We then used additional LMMs to consider whether the increase in proportion of time grooming was driven by a greater frequency (number of grooming interactions each individual was involved in divided by the time available for grooming at the refuge) or an increase in mean bout duration; both of these response variables were also log-transformed. We subsequently ran Wilcoxon signed-rank tests in SPSS 24 (as in *Observational data collection* and *Experiment 1 protocol*) to consider the grooming behaviour between specific categories of group members (see *Results*).

## Acknowledgements

We thank B. Rouwhorst and H. Yeates for access to their land, I. Carpenter, M. Layton, J. Linden, I. Shan, and N. Tegtman for invaluable support and assistance in the field, M. Aveling for the beautiful figure illustrations, and I. Braga-Goncalves, S. King and P. Kennedy for useful comments on the manuscript.

